# MerMADE: Coupled biophysical, eco-evolutionary modelling for predicting population dynamics, movement and dispersal evolution in the marine environment

**DOI:** 10.1101/2022.11.15.516611

**Authors:** Rebekka L Allgayer, Paul G Fernandes, Peter J Wright, Justin MJ Travis

**Affiliations:** Institute of Biological and Environmental Sciences, University of Aberdeen, Aberdeen AB24 2TZ, UK; The Lyell Centre, Heriot-Watt University, Research Avenue South, Edinburgh, EH14 4AP, UK; Marine Scotland Science, Aberdeen AB11 9DB, UK; Marine Ecology and Conservation Consultancy, Ellon AB41 8XY, UK

## Abstract

In order to understand patterns in species’ distributions, we need to understand the underlying mechanisms of dispersal, demography and evolutionary capability of these species. In the marine environment, few models combine these three key components likely due both to the computational challenges involved and the inherent challenges in data collection for parameterisation. To fill this gap, we have developed MerMADE, an individual-based, spatially explicit, eco-evolutionary coupled biophysical model for predicting population dynamics, dispersal and movement evolution in the marine environment (or aquatic environments in general). MerMADE combines dispersal in a 3D, hydrodynamically informed environment with population dynamics, demography and evolutionary functionality in order to investigate questions of connectivity, population persistence and evolution under environmental change and anthropogenic pressure. We illustrate its range of behavioural and physiological functionality using the lesser sandeel, *Ammodytes marinus*, as a case-study species in an invasion scenario. MerMADE’s flexibility in species-specific parameterisation makes it a widely applicable, exciting tool in future sustainable management and conservation of aquatic species under environmental change.

## Background

The underlying processes that shape species’ distributions have long fascinated ecologists with their unending complexity and mystery, with recent decades experiencing a surge in focus on how these are affected by environmental change (DeMarche Megan L. et al., 2019; Elsen et al., 2020; Román-Palacios & Wiens, 2020; Waldvogel et al., 2020). To avoid the common criticisms of correlative models, there have been significant advances in developing dynamic, mechanistic distribution models, especially in terrestrial environments (Zurell et al., 2022). However, for marine systems, while progress has been made to model movement with water currents (Christensen et al., 2018; Lett et al., 2008; North et al., 2008; Paris et al., 2013), there has not yet been development of models linking demography, dispersal and evolution to look at consequences of environmental change at large scales.

Progress has been made in the terrestrial sphere for coupling demography with dispersal in models developed to provide ecological forecasts. The RangeShifter platform (Bocedi et al., 2014, 2021) is a modelling framework for investigating species’ range dynamics, taking population dynamics, dispersal behaviour and evolution into account when predicting responses to environmental change. Similarly, STEPS (Visintin et al., 2020) combines the outputs of landscape models and species distribution models (SDMs) with population dynamics and dispersal, in order to predict population change across landscapes and after disturbance. However, these frameworks are rare in the marine environment, most likely due to the inherent difficulty in parameterising and quantifying dispersal and connectivity (Burgess et al., 2014; Carson et al., 2011; Swearer et al., 2019). The majority of benthic marine organisms utilise a planktonic larval stage (Bode et al., 2018; James et al., 2002; Swearer et al., 2019), making pelagic larval dispersal a major factor in determining connectivity between populations—often across wide distances—and therefore the viability of the greater metapopulation under predicted environmental change (Bode et al., 2019; Burgess et al., 2014; Mitarai et al., 2008). Hydrodynamic and oceanographic features of the marine environment play a major role in dispersal and resulting community structure and therefore also need to be accounted for mechanistically (Bertolo et al., 2012; Castonguay & Gilbert, 1995; Cowen & Sponaugle, 2009; Levine, 2003; Nathan et al., 2008; Sedell et al., 1989; Tonkin et al., 2014; Treml et al., 2008).

Recent years have seen an increase in the use of coupled biophysical models to study specifically the larval dispersal process and resulting connectivity (Bode et al., 2018; Miller, 2007). Several software packages are already available to determine connectivity metrics, with highly detailed hydrodynamic data. The Connectivity Modeling System (CMS), a multi-scale model that aims to generate connectivity and transitional probability matrices using tracked particle trajectories, is a released software that can be used for a range of focal species (Paris et al., 2013). Similarly, Ichthyop, developed by Lett et al (2008) is a publicly-available Lagrangian tool for modelling changes in ichthyoplankton dynamics in response to physical and biological factors (Lett et al., 2008). While these models can account for spawning times, they do not include population dynamics and do not allow for stage-structured, multi-generational contribution to the dispersing juvenile cohort. These models focus heavily on the “transfer” phase of dispersal, not focusing in detail on the “emigration” or “settlement” phases. Additionally, they do not allow for life history traits to evolve in response to environmental conditions (though Ichthyop does allow growth rate to be influenced by temperature (Lett et al., 2008)).

We therefore developed MerMADE, a model that incorporates population dynamics, a coupled biophysical, 3D movement model, and evolutionary functionality in order to ask larger-scale questions about the long-term effects of continued pressure. Due to our use of an Individual Based Modelling (IBM) framework, our model allows each individual to carry its own trait values, meaning we can apply our model to eco-evolutionary questions, allowing the changing environment to evolutionarily influence certain demographic or dispersal traits. We combine the effect of hydrodynamics with larval behaviour such as diel vertical migration and variable activity levels and flexible development based on growth rate parameters. Not only that, but these aspects are incorporated into a 3D representation of the aquatic environment, which is novel for a spatially realistic population dynamics simulator.

**Figure 1:**
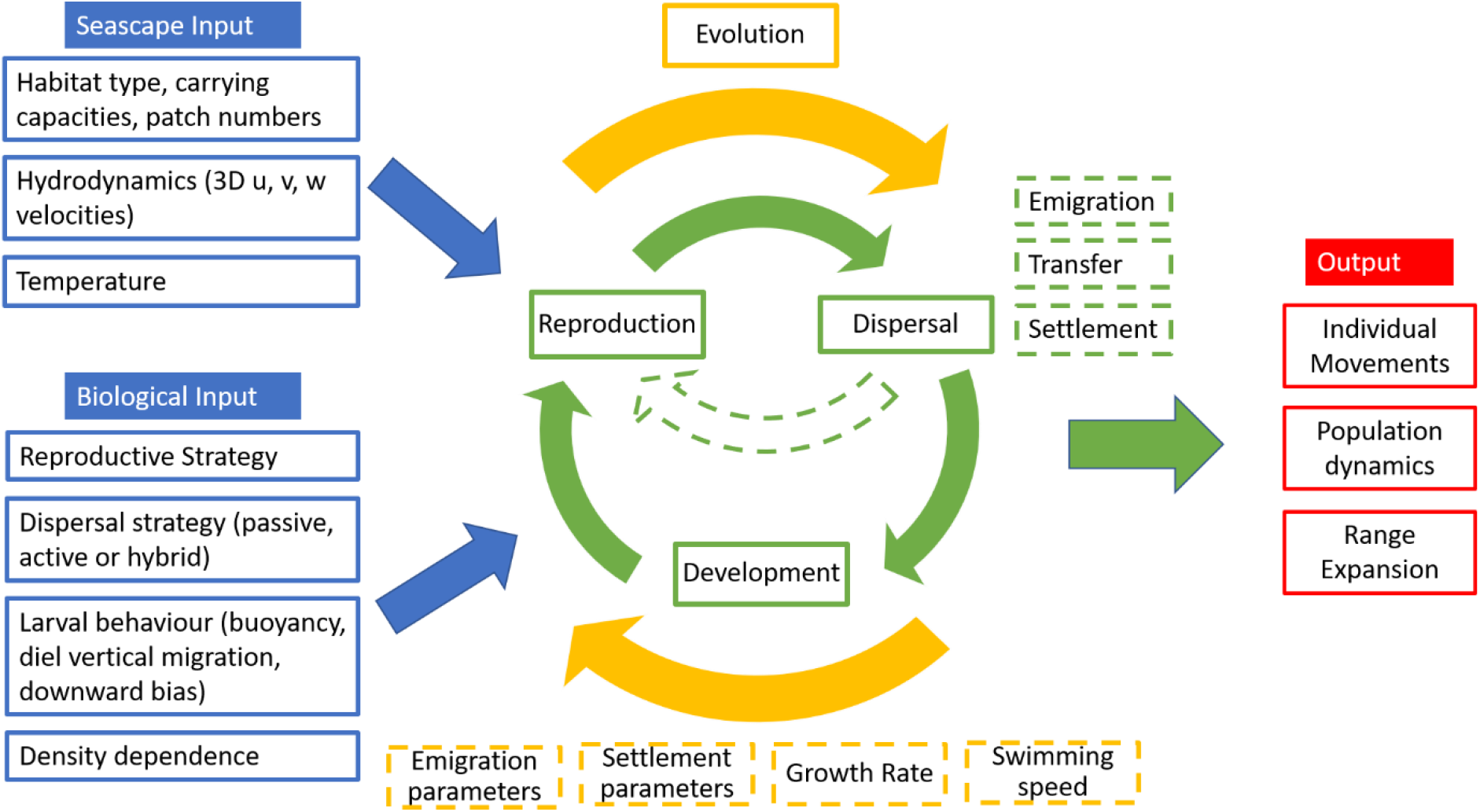
Schematic of how MerMADE is structured, showing various inputs and outputs as well as the dispersal and population dynamics modules incorporated into each annual cycle. Evolutionary capabilities carry on across generations.

By modeling the three phases of dispersal (emigration, transfer, and settlement) explicitly and allowing parameters in each phase to evolve, we can investigate drivers of distribution and species range under changing environmental conditions. Timings of behavioural changes during the transfer phase, for example, and temperature-dependent growth can affect which hydrodynamics individuals experience and therefore which suitable habitat they may have the ability to access, having knock-on effects on connectivity. Seeing the evolutionary effects of habitat fragmentation, climate change and fishing pressure between protected areas, for example, can give us information that will allow us to manage fish stocks sustainably and protect vulnerable or threatened species effectively in a changing world.

## Methods and Features

### Dispersal

This model is built on the foundations of the eco-evolutionary modelling framework RangeShifter, and therefore incorporates much of the same functionality in terms of population dynamics and the three-stage dispersal process (please refer to Bocedi et al. 2014 and 2021 for more detail). Where the main novelty lies in MerMADE’s case is during the transfer phase, the movement from an individual’s natal patch to its chosen site for reproduction.

As each individual moves, it encounters cells of varying habitat types (ie water, suitable or unsuitable habitat). Hydrodynamic information is stored as velocity values (u (eastward velocity), v (northward velocity) and w (upward velocity) in a Cartesian vector format), which are then used in the following equation to calculate the angle and magnitude of the current vector. Since this is a 3D vector, two angles are needed to describe its direction.

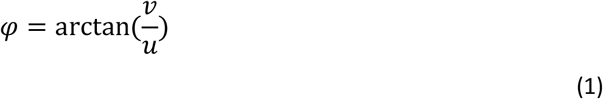

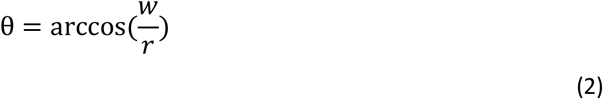

Where *r* is the magnitude of the 3D vector, which also represents the current’s speed in that cell, and is calculated by:

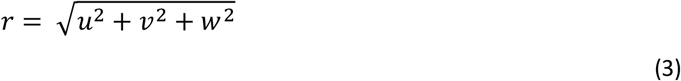

Angles are handled in radians (0, 2π).

We now know in which direction the current is flowing and how quickly, but the dispersal method of the individual will determine how it reacts to this information. Passive dispersers will be highly influenced by the current while active dispersers may be able to make additional decisions that will dictate in which directions they swim. The calculated *φ* angle is then used as a mean in a circular Wrapped Cauchy Distribution to introduce stochasticity to the model. A concentration parameter *ρ* is used to indicate how closely sampling should adhere to the given mean of the distribution. For passive individuals, *ρ* = 0.9, but the *ρ* for active dispersers is user-defined (0 — 0.9). A value *ρ* = 0 would indicate an entirely active disperser that is not influenced at all by current and increasing *ρ* represents increased attention to current directionality during decision-making.

Using *φ* and *θ*, the coordinates of the new position can then be calculated using:

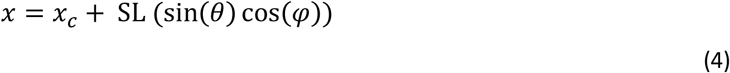

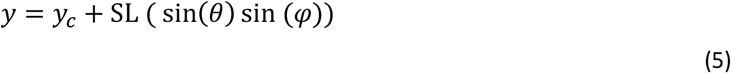

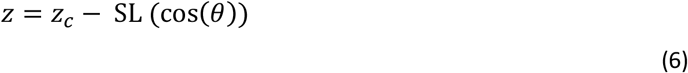

where *x_c_, y_c_, z_c_* are the coordinates of the current position (in metres), and SL is the step length or swimming speed of the individual in m/hr. If the individual is passive, SL is the speed of the current, as they have no swimming power of their own. Unless an individual has a very high swimming speed or the spatial resolution is very fine, movements on a temporal scale finer than hours is unlikely to be highly significant.

Active individuals are also able to store past movements in memory, thereby increasing their directional persistence, avoiding overly laborious and inefficient movements. Additionally, even if individuals have swimming capabilities, they still experience the proportionate effect of the current on their movements. For example, if they go against the current, they will only be transported the appropriate fraction of their SL forward, as the current will have acted in the opposite direction. Similarly, if going with the current, they experience the equivalent of a “tailwind” and are able to travel further. Therefore, we take a weighted average of the two vectors: the direction towards suitable habitat and direction of current flow (Equations 7-10), using Equations 4-6.

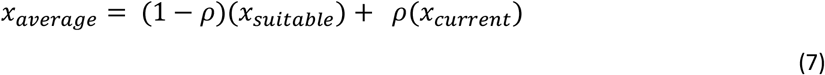

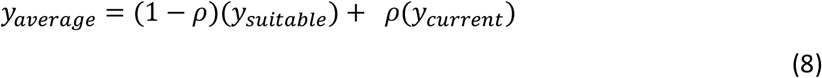

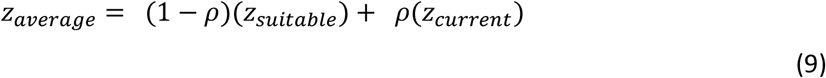

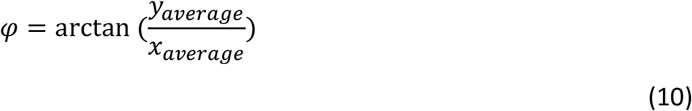

Many organisms that use pelagic larval dispersal have both a passive and an active phase. This means that after a developmental period, the juvenile is then morphologically competent enough to be able to influence its movements and potentially detect and move towards suitable habitat. The model also allows this functionality, in that after a user-defined competency period is reached, a passively dispersing individual is assigned a swimming speed and searches the environment for suitable habitat. This also gives it the ability to assess a habitat patch or cell when it comes across one and resuspend and continue searching, should it decide not to settle. Detection distance is user-defined in number of cells (making it resolution dependent), and can represent detection via sight, sound or olfactory cues. The ability of the individual to follow these cues and move towards suitable habitat depends on its dependence on the current, *ρ*.

Alternatively, or perhaps additionally, individuals may undergo larval behaviour such as diel vertical migration (dvm) which affects their vertical distribution in relation to day-and-night cycles but does not give them the ability to influence their horizontal movement. A user-defined dvm range is used to determine upper and lower boundaries, and positions within the water column are then sampled within these restrictions every 12 hours to simulate day vs. night, and we assume juvenile release occurs in the evening. We acknowledge that this is simplified and that, depending on time of year and geographical location, an even day-night split may not be accurate. Improvements to the accuracy of dvm within this model is an area for future development.

MerMADE therefore allows for four stages of dispersal activity: completely passive, passive with diel vertical migration, active pre-competency, and active with settlement competence. The user can choose any combination of these activity levels, given suitable information on minimum times and/or sizes for when to switch to the next level.

The timing of when to increase in activity level can be time- or size-dependent. For example, if behaviour is time-dependent, the user could specify that individuals begin to exhibit dvm after 3 days dispersing. Alternatively, this decision could be made more mechanistically and be size-dependent, so the switch only happens after an individual is, say, 5mm in length. The user can choose between two methods of calculating an individual’s length: linear,

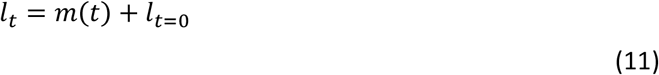

where m is the slope of the linear relationship; or a modified Gompertz curve (Régnier et al., in prep)

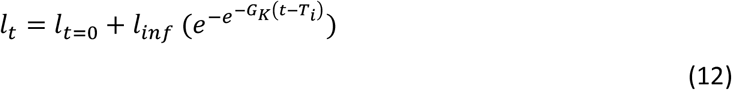

where *l*_*t*=0_ is the size at hatching, *l_inf_* is the maximum length an individual can reach, where this function reaches an asymptote, *G_κ_* is a growth parameter, *t* represents the time left for dispersal until PLD is reached, and *T_i_* is the inflection point or earliest date of settlement. The modified Gompertz from Régnier et al. 2022 (in prep) was chosen because it takes into account size at birth and can therefore play a role in the timing of behavioural changes. For the linear method of calculating size, the user can include temperature-dependence, if, for example, the environmental change being investigated is change in water temperature. Growth only occurs during dispersal, so the model stops tracking an individual’s size after settlement.

Settlement of passive individuals (where there is no active dispersal phase at all) requires some assumptions to be made. In some cases, larvae “settle out” of the water column but are still considered passive and don’t have a swimming speed which means they are at the mercy of local currents and eddies as deciding factors between settlement or not. Depending on the resolution of the rest of the seascape, these fine scale interactions may not be captured. A settlement zone option is therefore available as a function in MerMADE to define a distance from suitable habitat within which an individual is simply considered to have settled, regardless of activity level. For actively dispersing individuals, settlement is completely dependent on their ability to seek out and swim toward suitable habitat.

### Evolution

The absence of evolutionary functionality in most models predicting species’ responses to environmental change implicitly makes the assumption that traits are fixed and species can’t adapt to their new conditions. However, this rules out a potential coping mechanism, making conclusions drawn about the capability of a species to persist locally, or regionally, under environmental change incomplete. MerMADE therefore includes evolutionary functionality in several traits: emigration probability, age/size at activity and competency levels, growth rate and settlement probability. Each of these traits is allowed to evolve given a mutation rate and size in response to given environmental conditions. This mutation occurs when individuals inherit their parent(s)’ genetic information, with individuals storing the mutated genotype, ready to pass it onto its own offspring.

Having traits able to evolve in all three stages of dispersal allows us to tease apart crucial factors that decide the success of dispersal, as well as the influence that other physiological processes might have.

## Example Application

MerMADE is relevant for a wide range of species due to its flexibility in behavioural modes during dispersal as well as its ability to incorporate physiological components such as larval growth and an enforced competency threshold. To demonstrate this flexibility, we applied MerMADE to a hypothetical invasion (or reintroduction) example. We use this example to illustrate how different dispersal behaviours can be incorporated in the model as well as larval growth and associated changes in swimming competency. To provide the example, we use the physiological characteristics and life history traits of the lesser sandeel, *Ammodytes marinus*, a species which has a patchy distribution on sandbanks in the North Sea. The model choices and key parameter values used in the different simulations illustrated in Figure 2 are summarised in Table 1 while the table of parameter values we use can be found in the Supplementary materials. The results demonstrate very clearly that the interplay between species-specific physiology and behaviour has substantial effects on the spread of an invasive species—or, for that matter, the success of a reintroduced species (Figure 2).

**Table 1:** MerMADE simulations run to illustrate the effect of incorporating behaviour and physiology (growth and competency) into potential spread of invasive species

| Simulation | Passive phase | Diel vertical migration | Active phase | Competency: | Growth? |
| --- | --- | --- | --- | --- | --- |
| Behaviour Only | ✓ | - | - | Day 0 | - |
|  | ✓ | Day 20 | Day 45 | Day 0 | - |
|  | - | - | Day 0 | Day 0 | - |
| With Growth | ✓ | - | - | 5.33mm | ✓ |
|  | ✓ | 10mm | 26mm | 5.33mm | ✓ |
|  | - | - | 5.33mm | 5.33mm | ✓ |
| Growth & Delayed Competency | ✓ | - | - | 26mm | ✓ |
|  | ✓ | 10mm | 26mm | 26mm | ✓ |
|  | - | - | 5.33 | 26mm | ✓ |

In the top row of Figure 2, behaviour is the driving factor between the different scenarios. Even though most passive individuals are caught in a settlement buffer radius around their natal patch since competency is not a limiting factor, those that do disperse are carried further on the current until they successfully settle elsewhere. The passive behavioural mode therefore has a greater spread when compared with hybrid dispersers, who, after their passive phase, settled on the nearest patches. Population size therefore booms on the original site, which we do not see to the same degree in the hybrid or active scenarios. Spread is very limited in the hybrid mode due to the hydrodynamics keeping individuals in a cyclical movement pattern during their passive phase, favouring local retention if the model allowed it (active dispersers are assumed not to choose to settle on their natal patch). The trajectory of spread would most likely continue north out of the extent of the map over longer periods of time since the most north-easterly patch has been colonised. The immediately active behavioural mode is not slowed by the passive phase and can therefore make the most of the available habitat, reaching the maximum extent within the 50 years.

**Figure 2:**
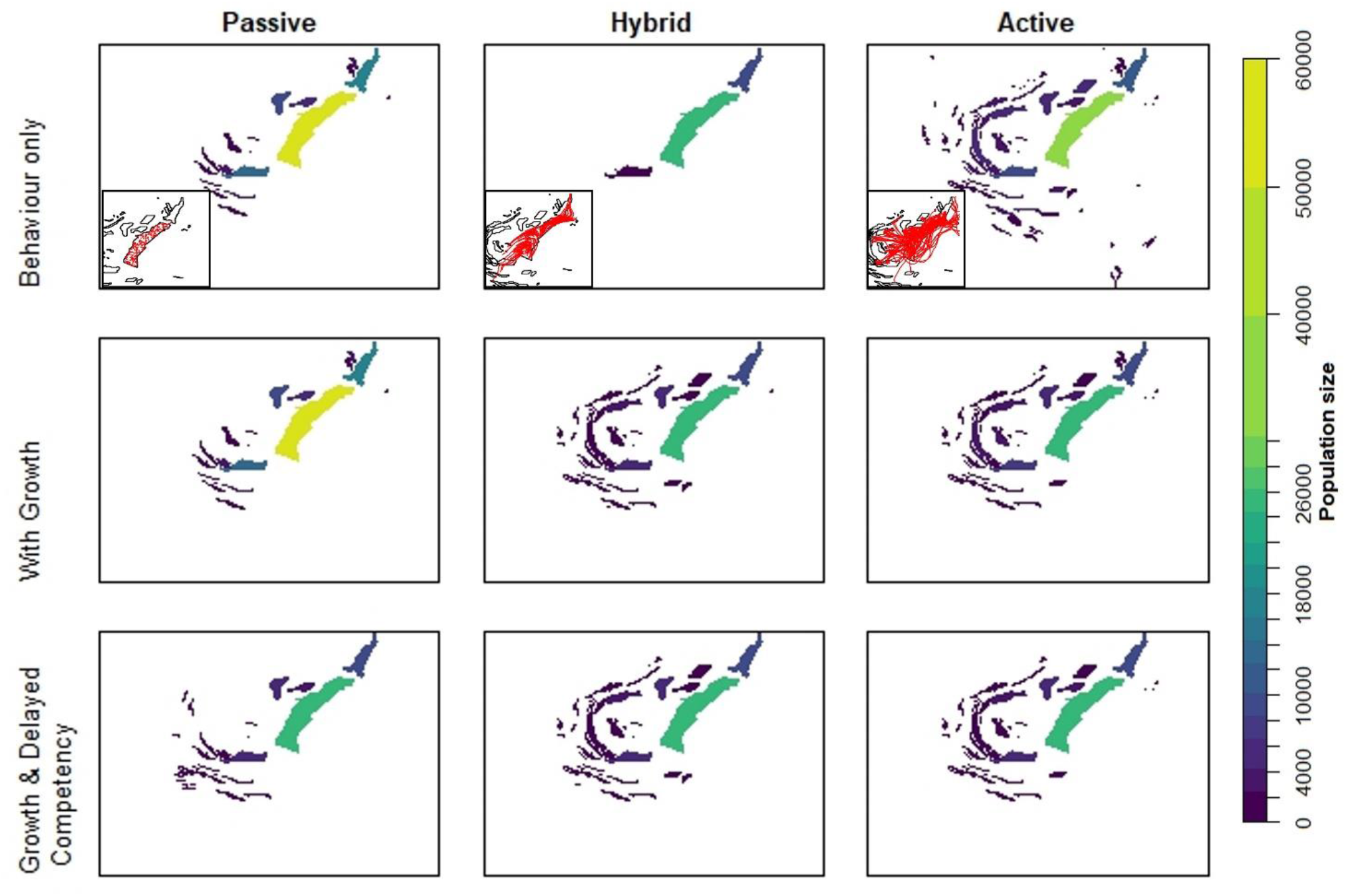
Results of an invasion scenario after 50 years using *Ammodytes marinus* as a case study species. Simulation details can be found in Table 1. Colours indicate population size at each patch. Inserts in the top row show differences in movement tracks (lines shown in red) due to complexity of behaviour during dispersal.

The addition of growth rate only truly effects the hybrid and active modes, as swimming speed and therefore ability to combat the current is dependent on body size. For the hybrid dispersers, the switch between diel vertical migration and active dispersal is also size dependent. The variable swimming speed enables individuals to settle successfully on more neighbouring patches, thereby driving the invasion over the 50-year simulation. The period of time at lower swimming speeds for smaller active individuals compared to the simulation that did not incorporate growth restricts the invasion capabilities of the species.

The overall result from a presence-absence perspective when incorporating delayed competency into the model is not that drastic for hybrid and active dispersers by the end of the simulation. However, the population sizes of the passive individuals change significantly, as delayed competency hinders settlement in the natal and neighbouring patches. Information on presence as well as abundance is important to consider for population dynamics and connectivity predictions going forward in an invasion (or reintroduction) scenario.

This example simultaneously illustrates the range of traits MerMADE can accommodate and the importance of considering behavioural and physiological constraints on dispersal.

## Conclusion

The inherent difficulties in quantifying dispersal and connectivity empirically in the marine environment (Carson et al., 2011; Swearer et al., 2019) have impeded progress in mechanistic modelling practices. This leaves questions of metapopulation persistence in applications such as marine protected area network design considering functional connectivity, but not realised connectivity (Burgess et al., 2014; Figueira, 2009). Environmental change can present itself in many forms: temperature and salinity fluctuations, more frequent extreme weather events, habitat fragmentation or reduction in habitat quality, increased mortality due to resource harvesting such as fisheries, among others. Changes such as these exert a selective pressure on certain dispersal as well as demographic traits to adapt to new conditions or can cause the species’ range to shift to more suitable habitat. With either of these responses, it is important to take them into account when designing future management strategies so that precious time, effort and financial resources are not wasted on actions that are not “future proof”.

Though many studies have focused on creating very detailed larval transport and connectivity models (Bode et al., 2019; Lett et al., 2008; North et al., 2008; Paris et al., 2013), not many have combined these with demography (Burgess et al., 2014; Carson et al., 2011; Figueira, 2009), and even fewer, if any, incorporate evolutionary potential. We developed MerMADE to address the lack of these frameworks in the marine environment. MerMADE has the ability to track population-scale changes over time, incorporates species-specific behavioural and physiological components in the dispersal process, and allows the species to adapt to environmental change through mutation and inheritance of beneficial genotypes. This coupling of multiple mechanistic components makes MerMADE uniquely flexible and able to be applied to a range of applications and research questions.

While we believe the sacrifices in detail that we have made in the areas of hydrodynamics and buoyancy are reasonable and acceptable, we do acknowledge that they are important considerations and will work to further develop MerMADE to minimise this effect. However, the flexibility in the transfer phase of dispersal as well as the incorporation of population dynamics and evolutionary capability makes MerMADE uniquely suited to addressing questions of population connectivity and viability under changing conditions, natural or anthropogenic. Future-proofing management actions is becoming increasingly important as climate change and harvest of marine resources become more extreme. Any knowledge that can aid in understanding movement patterns and potential species’ responses to such stressors will be invaluable in our attempts to minimise negative impacts on the marine environment.

