## Supplemental Table 1 for "MerMADE: Coupled biophysical, eco-evolutionary modelling for predicting population dynamics, movement and dispersal evolution in the marine environment"

Table S.1: Parameter values and references for A. marinus used in MerMADE simulations. It is worth noting that while Step Length (SL) is given here in BL/s, MerMADE multiplies this out to a unit of m/hr, depending on body size at any given timestep.

| | Parameter | Value | Reference (if applicable) |
|---|---|---|---|
| | Maximum age | 10 years | |
| Emigration | Emigration Probability | 0.8 | |
| Transfer | Pelagic Larval Duration (PLD) | 70 days | Within the range from Wright and Bailey 1996; Régnier et al. 2017 |
| | Buoyancy Range | 0-80m | |
| | Diel Vertical Migration (dvm) range | 10m (0-10m at night, 70-80m in the day, unless seafloor is shallower) | Jensen et al., 2003 |
| | Minimum size at dvm | 10mm | Yamashita et al., 1985; Jensen et al., 2003 |
| | Minimum size at active | 26mm | Régnier et al 2021 (in prep) |
| | Step Length (SL) when active | 1 BL/s | |
| | Dispersal-related mortality | 0.042/day | Régnier et al. 2017 |
| Growth (modified Gompertz ) | $l_0$ | 5.33 mm | Régnier et al. 2021 (in prep) |
| | $l_{inf}$ | 67.04 mm | Régnier et al. 2021 (in prep) |
| | $K$ | 0.03696889 | Régnier et al. 2021 (in prep) |
| | $T_i$ | 53 days | Régnier et al. 2021 (in prep) |
| Settlement | Minimum size at competency | 26mm | Régnier et al. 2021 (in prep) |
| | Settlement Probability | 1 | |
| | Settlement buffer | 4.5km | |
